# The Functional effects of voluntary and involuntary visual phantom color on conscious awareness

**DOI:** 10.1101/286716

**Authors:** Shuai Chang, Joel Pearson

## Abstract

The constructive nature of vision is perhaps most evident during hallucinations, mental imagery, synesthesia, perceptual filling-in, and many illusions in which conscious visual experience does not overtly correspond to retinal stimulation: phantom vision. However, the relationship between voluntary and involuntary phantom vision remains largely unknown. Here, we investigated two forms of visual phantom color, neon phantom color spreading and voluntary color mental imagery and their effect on subsequent binocular rivalry perception. Passively viewing neon phantom color induced time sensitive, suppressive effects on spatially non-overlapping subsequent binocular rivalry. These effects could be attenuated by rotating the color-inducers, or like color imagery, by concurrent uniform luminance stimulation. The degree of neon color induced rivalry suppression predicted the degree of voluntary color imagery facilitation, both on subsequent rivalry perception. Further, these suppressive and facilitative effects were additive when experienced successively. Our results suggest potential sensory mechanistic commonalities between voluntary and involuntary phantom vision.

## Introduction

The constructive nature of normal human vision can be illustrated by many examples of perceptual completion in which the brain generates or fills in missing visual features that are absent from retinal stimulation. Such visual phantoms or non-retinal vision can be dynamic or static, depending on the particular induction scenario (Pearson & Westbrook, 2015). This non-retinal visual experience has been called phantom vision. Interestingly, non-retinal phantom sensory experience can be generated *voluntarily* by most individuals in the form of mental imagery that can be formed in the absence or presence of incoming retinal stimulation (Pearson, Naselaris, Holmes, & Kosslyn, 2015). In contrast, phantom vision can also be formed involuntarily, and two striking examples of such visual phantoms include phantom moving gratings and the neon color spreading illusion (Meng, Remus, & Tong, 2005; Sasaki & Watanabe, 2004).

The neon color spreading illusion can be induced by four groups of concentric circles, when the inside segments of the circles share a common color and the edges are aligned with the contour of a ‘missing’ shape, conscious experience of the color typically fills into the blank inner area, creating the experience of color where there is no retinal stimulation (Figure 1A). This type of visual experience does not directly originate from retinal input, but is constructed by the brain, based on the inducer information, hence referred to as non-retinal or phantom color (Pearson & Westbrook, 2015; Sasaki & Watanabe, 2004).

**Figure 1.**
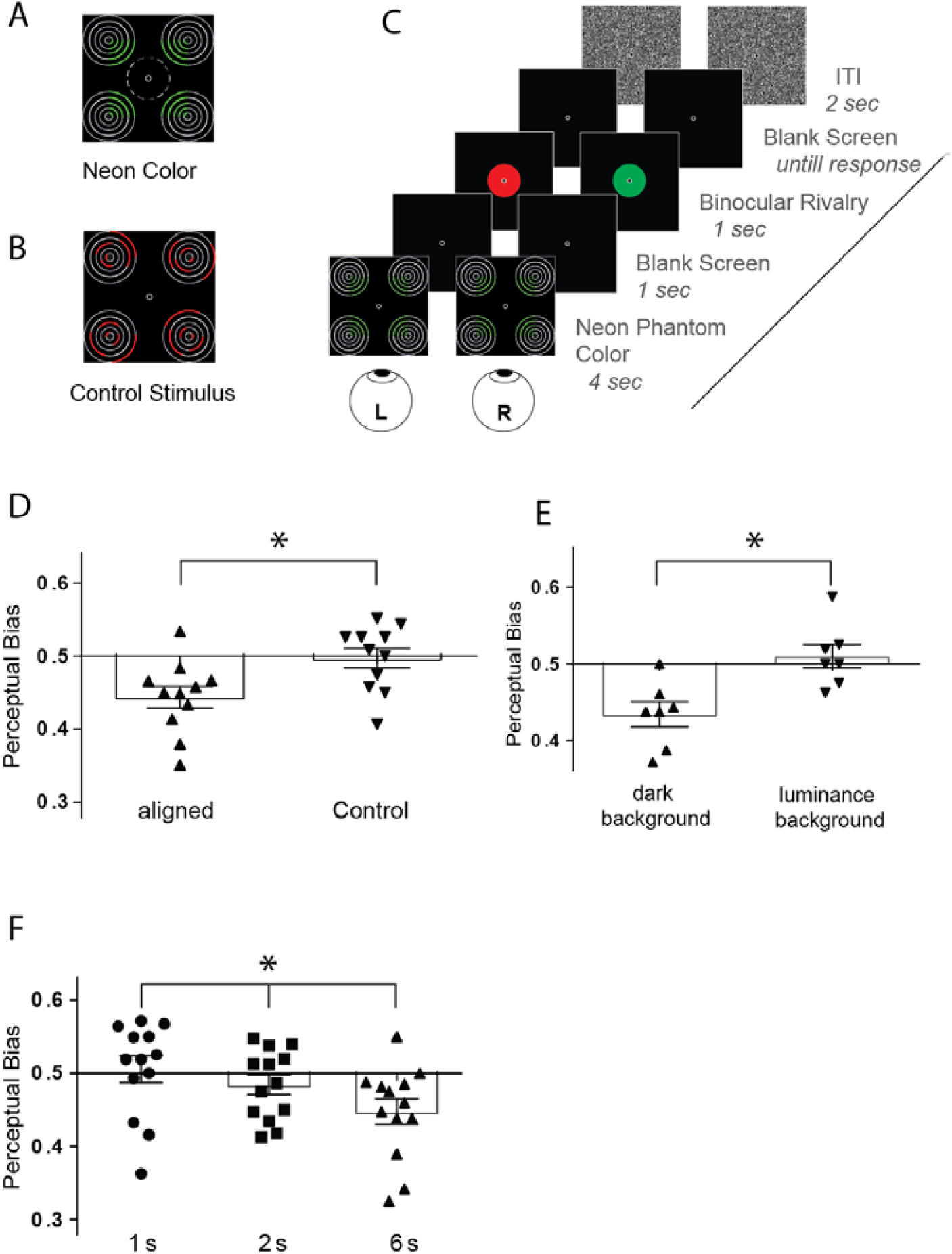
A: Neon color spreading illusion stimuli used in the study. The circle in dashed line at the center indicates the location of binocular rivalry stimuli and was not shown in the actual stimulus. B: An example of stimuli in random condition of Experiment 1. C: Trial timeline in Experiment 1. D: Results from Experiment 1. The mean perceptual bias in aligned condition was M=0.444 and it was significantly different from chance. In the random condition mean perceptual bias M=0.497 was not significantly different from chance. E: Results of Experiment 2. The mean perceptual bias in the dark background condition was M=0.443. In background luminance condition, the mean perceptual bias was not significant from chance level M=0.510. F. Results from Experiment 3. Significant difference from the three presentation conditions. Error bars show ±SEM. * p<.05

**Figure 2.**
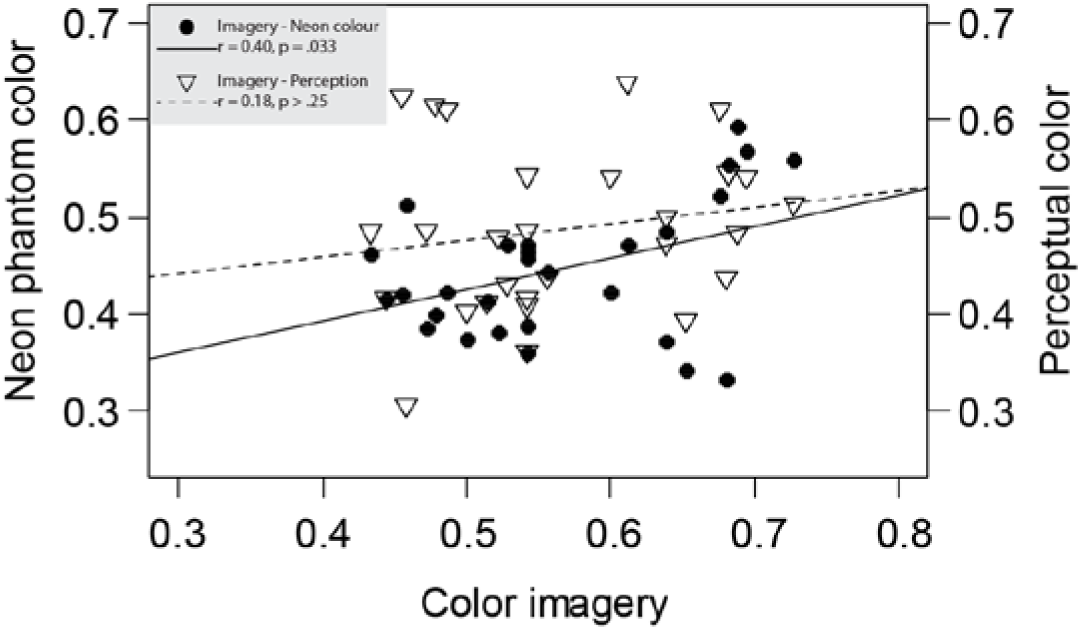
Significant correlation was found between the functional effects of voluntary imagery and neon phantom color (r = 0.40, p = .033, shown as solid dots and solid line). Effects of voluntary imagery and perceptual stimuli were not significantly correlated, shown as empty triangle and dash line.

Neuroimaging studies have shown that such involuntary phantom vision is related to enhanced BOLD response in primary visual cortex (Meng et al., 2005; Sasaki & Watanabe, 2004). Meng et al. (2005) reported enhanced BOLD signal in regions of V1 corresponding to the moving phantom pattern. Likewise, research has also found similar V1 activation for both involuntary neon phantom color and its matched perceptual color (Sasaki & Watanabe, 2004).

Seeing that both forms of phantom vision are examples of the constructive nature of the visual system, a dichotomous framework for phantom vision, analogous to the sub-types of attention: endogenous and exogenous, has been proposed (Pearson & Westbrook, 2015). This proposal suggests the two types of phantom vision share a common sensory substrate in sensory cortex. However, research directly comparing the two forms of phantom vision is largely non-existent. Much of the difficulty of examining phantom vision stems from its inherently private and subjective nature. However, recent behavioral work on voluntary phantom vision has developed reliable and indirect methods of investigation using binocular rivalry (Pearson, 2014). Here we adapt this rivalry-method to assess involuntary phantom color in the form of neon color spreading. Using the one indirect perceptual method, color rivalry, we measure and compare the sensory strength of both voluntary and involuntary phantom color.

Experiment 1 to 3 tested the functional effects of neon phantom color on subsequent color rivalry and how the inducing pattern structure, uniform background luminance and pattern presentation duration influence the effect. Experiment 4 measured the sensory strength of mental imagery, neon phantom color and retinal color perception, and mental imagery, and found neon phantom color were correlated. Experiment 5 showed the effects of consecutive imagery and neon color interact with each other and are additive. This additive effect was further examined in Experiment 6. These results provide novel behavioral support for the hypothesis that these two forms of constructive non-retinal vision rely on common mechanisms in the visual processing hierarchy (Pearson & Westbrook, 2015).

## General Method

In the current study, we used the binocular rivalry paradigm to investigate the functional effects of phantom vision. This method has been used in previous studies on voluntary mental imagery, and found that after forming a mental image of one of two patterns, the perception of a brief binocular rivalry stimulus can be biased towards seeing the prior imagined pattern (Pearson, Clifford, & Tong, 2008). By calculating the perceptual bias on the briefly presented rivalry stimulus, the strength of the prior sensory representation can be quantified (Pearson, 2014). This indirect quantitative method has been used in voluntary imagery studies. Previous studies have found that the priming or bias effect is specific to the spatial orientation (Pearson et al., 2008) and the retinotopic location of the imagined pattern (Chang, Lewis, & Pearson, 2013), is attenuated by concurrent background luminance (Pearson et al., 2008; Sherwood & Pearson, 2010), can be differentiated from visual attention (Pearson et al., 2008), is related to subjective reports of vividness (Pearson, Rademaker, & Tong, 2011) and is reliable over a two week re-test (Bergmann, Genç, Kohler, Singer, & Pearson, 2015).

The detailed procedures of each experiment will be provided in the following experimental sections.

### Participants

The final number of participants in Experiments 1 to 6 were 13, 9, 14, 54, 11 and 20, respectively, the range of these numbers were based on prior work looking at mental imagery and visual priming (Chang et al., 2013; Keogh & Pearson, 2011, 2014; Pearson et al., 2008). Data from some participants was excluded from analysis, and the exclusion criteria are described in Experiment 1. All participants had normal or corrected-to-normal vision and provided written informed consent. They were rewarded with $15 per hour for their participation. All experiments were approved by UNSW Human Research Ethics Advisory Panel for Psychology.

### Apparatus and stimuli

In all experiments, stimuli were displayed on a DELL Trinitron P1130 CRT monitor with 1280 × 1024 resolution and an 85-Hz refresh rate. Stimuli were generated using Matlab 8.0.0 (R2012b) and the Psychophysics toolbox (Brainard, 1997; Pelli, 1997), running on a PC with an AMD Radeon HD7700 graphics chip. In all experiments, stimuli were displayed within a square area with a side length of 9.3 degrees and a circular bulls-eye fixation mark (diameter = 0.3° visual angle) was used to help participants maintain fixation. During the experiments, participants were seated in a darkened room with their heads stabilized using a mirror stereoscope and chin rest.

The neon color spreading stimulus contained four Pac-men-like patterns and each pattern was formed by five concentric circles. The colored parts of the concentric circles were rendered in red or green (See Figure 1A). Binocular rivalry stimuli consisted of two disks of red and green with the diameter of 3°. The CIE color values of the circles were: red: x = 0.61, y = 0.37; green: x = 0.28, y = 0.63. Importantly, the neon phantom color inducers made from the concentric circles and the binocular rivalry stimuli *did not* spatially overlap (as illustrated in Figure 1A).

In Experiment 1, we also ran a control condition, the color arcs in the concentric circles were rotated by random angles around their circle centers in each trial, so that there was no alignment among the color inducers (an example is shown in Figure 1B). The stimuli in the aligned and random control conditions varied in the orientations of the color arcs but the total number of colored pixels did not change.

In Experiment 4, perceptual stimuli of red and green disks were included, and the size and CIE values of red and green were identical to those in color binocular rivalry stimuli. Luminance of the perceptual stimuli were 40% of the values in the binocular rivalry stimuli.

Rivalry catch trials were included to probe for potential non-perceptual bias in each experiment. Catch trials were created by combining the two colors together, where half was red and the other half was green. This catch trial stimulus was presented simultaneously to both eyes, ensuring that the two colors were perfectly balanced and no actual rivalry could occur. Participants were required to report the more dominant color between the two rivalry colors in all test trials, so when they were presented with this kind of catch trial they should respond by pressing the “equal (mixed)” key. This type of catch-trial allows us to ensure participants’ responses to dominance were based on their perception and not any decisional bias. If participants report rivalry dominance based on some non-perceptual bias, then significant bias away from chance/veridical would be observed in these catch trials.

### Data and code availability

The data and custom codes that support the findings of this study are available from the corresponding author on reasonable request.

## Experiment 1: Functional effect of neon phantom color

### Participants

13 participants were recruited for this experiment, data from 2 participants were excluded from analysis because they had high proportions of ‘mix-response’ (greater than 10% of trials). For the remaining data, trials with a ‘mix-response’ were excluded from further analysis.

### Procedure

An eye-dominance test was first conducted to balance the dominance of red and green in the color rivalry (Pearson, 2014). Relative luminance of the two color patches was adjusted during this test in order to minimize potential eye bias. For the detailed procedures of eye-dominance test, please refer to Pearson et al. (2008) and Pearson (2014).

In the aligned main condition, at the beginning of each trial, participants were played a beep, and then observed neon phantom color for 4s of either red or green (randomly shuffled). After 1s blank screen, they were presented with red-green binocular rivalry stimulus for 1s. They then reported the dominant color via a key press where key “1” for red dominance, key “3” for green dominance and the middle “2” for equal dominance of the two colors. Followed by 2s of dynamic white noise presented between successive trials. (See Figure 1C). In the control condition, the neon phantom color inducer was replaced by the controlled illusory inducer as described above.

Each participant completed 3 blocks, and each block included 40 trials and 8 catch trials. The number of neon phantom color and control trials was identical in each block and they were presented in random order.

### Results

Data from Experiment 1 is shown in Figure 1D. We found a significant suppressive effect of the neon phantom color on subsequent binocular rivalry perception in the main aligned condition (M = 0.444, t(10) = 3.75, *p* = .004, Cohen’s d = 2.37), with no significant bias in the control condition (M = 0.497, t(10) = 0.206, *p* > .25, Cohen’s d = 0.13). There was significant difference between the two conditions, t(11) = 2.57, *p* = .028, Cohen’s d = 1.11). We also included catch trials with fake non-ambiguous rivalry stimuli that are not overtly susceptible to prior perceptual effects, to test for any non-perceptual criteria or decisional bias. The mean non-perceptual bias in the catch trials was 49.6%, which was not significantly different from chance level (*p* = .167).

These results suggest that neon phantom color induces a functionally suppressive effect on subsequent conscious perception, e.g., when you see the phantom neon green, you tend to see red as dominant in subsequent rivalry. In the control condition, the color arcs in the illusion inducers were rotated each by a random amount (Figure 1B), which breaks the generation of illusory color, while keeping the overall amount of perceptual color unchanged. Under this condition, no significant effects were found, indicating that the bias on subsequent awareness was not due to the presence of colored pixels in the inducers, but likely due to the illusion formed by the aligned orientation.

## Experiment 2: Disrupting phantom color with sensory stimulation

### Participants

9 participants were recruited for this experiment, data from 2 were excluded from analysis because of a high proportion of mixed percept responses (more than 10% of main trials). For the remaining data, trials with ‘mix-response’ were excluded from analysis.

### Procedure

Experiment 2 procedure was similar to Experiment 1, except that in the luminance background condition, a disk of uniform white was presented at the center of the neon phantom color for the entire 4s. The white disk had no overlap with any part of the inducers, but did cover the space where the subsequent color rivalry stimulus was presented. Each test block included 40 trials and 8 catch trials. Each participant completed 4 blocks of test.

### Results

Data from Experiment 2 is shown in Figure 1E. When neon phantom color was induced on a dark background, it again had a suppressive effect on subsequent binocular rivalry (M = 0.434, t(6) = 4.043, *p* = .007, Cohen’s d = 3.30), replicating our original finding. However, when presented with a white background, the effect was not significantly different from chance (M = 0.510, t(9) = 0.639, *p* > .25, Cohen’s d = 0.522). We also found significant difference between the two conditions (t(6) = 2.554, *p* = .043, Cohen’s d = 1.81). Again, no significant bias was found in catch trials (M = 0.5, *p* = 1). These results indicate that the neon phantom color, just like voluntary color imagery, can be disrupted by concurrent afferent, retinal stimulation.

## Experiment 3: Temporal dynamics of neon phantom color

### Participants

14 participants were recruited in this experiment and data from 1 participant was excluded because of their high proportion of mixed rivalry responses (>10% of trials). Mixed responses were excluded from analysis.

### Procedure

The procedure of Experiment 3 was same as Experiment 1, except for that the duration of phantom color presentation was varied. There were three duration conditions: 1s, 2s and 6s. The duration of all trials in one testing block were the same. There were 48 trials in each test block including 8 catch trials, and participants were required to complete 2 blocks of each type.

### Results

Figure 1F shows the degree of suppression for the three different phantom color durations. A repeated measures ANOVA found a main effect of duration (F(2,12) = 4.60, *p* = .025, η^2^ = .0.15). Post-hoc tests showed that there was a significant difference between the condition of 1s and 6s presentation duration (adjusted *p* = .01). Again, the average non-perceptual bias in catch trials was 49.0%, which was not significant different from chance (*p*=.153). These results show that longer color induction lead to stronger suppressive effects on the subsequent binocular rivalry perception.

## Experiment 4: Comparing voluntary and involuntary phantom color

### Participants

54 participants were recruited in this experiment. Data from 22 participants were excluded, 13 of these were excluded because of their high proportion (> 10%) of equal rivalry responses, 1 because of strong bias in catch trials, which was at 0.854, and 8 due to a high proportion of responses to one color (> 90%; e.g. almost always just reporting red) in at least one of the three conditions. These extreme results indicate that they may have not attended or performed the experiment appropriately, or had an imbalance in their rivalry eye dominance (Pearson et al., 2008). After this exclusion stage, data from 4 participants was found to exceed 2 standard deviations, and they were excluded from further analysis because outliers can significantly influence statistic tests (Osbourne & Overbay, 2004; Zimmerman, 1994). After this, 28 participants remained with these criteria.

### Procedure

The procedure for neon phantom color was same as the dark background condition in Experiment 1.

For the mental imagery test, participants were presented with a letter cue for 1s to indicate which color to imagine for the following 4s. Following this, color binocular rivalry was presented for 1s, and participants pressed the corresponding key to indicate the dominant color or mixed percept.

For the perceptual color test, the procedure was identical to that for imagery, except instead of the letter cue and voluntary imagination task, participants passively viewed a weak, low luminance color patch. The perceptual stimuli were set at 30% of the rivalry stimuli luminance.

In all three conditions, there were 48 trials in each test block including 8 catch trials. The number of red and green color trials was the same in each block and they were randomly intermixed. Participants completed 2 blocks of each test.

### Results

There was a significant correlation between perceptual biases produced by color imagery and neon phantom color (r = .40, *p* = .033), data shown in figure 2. The correlations between the effects of neon phantom color and perception (r = .27, *p* = .16) and color imagery and color perception (r = .18, *p* > .25), were not significant. There was no significant difference between color imagery vs. neon phantom color, and the color imagery vs. color perception correlations (z-score = 0.977, *p* = .328) (Lee & Preacher, 2013). The average decisional bias in catch trials in the three experiments were between 49.8% and 50.1%, and none were significantly different from chance level (*p*s > .3). In addition, perceptual biases in binocular rivalry trials were not significantly correlated with decisional biases in the catch trials (*p*s > .1)

Binocular rivalry was the common depended measure across all these three conditions. However, we did not see significant correlations between all conditions. This suggests that the correlation between imagery and neon color was not entirely due to the common dependent measure of rivalry. Given this, the positive correlation between voluntary and involuntary phantom color suggests some shared neural mechanism between voluntary and involuntary phantom color.

## Experiment 5: Joint effect of voluntary and involuntary phantom color

The above experiments showed commonalities between the functional effects of neon color and color imagery, suggesting potential shared mechanisms between the two forms of phantom vision, and this could be related to the activation of V1. In the Experiment 5, we tested the joint effects of voluntary and involuntary color. If the two processes have shared mechanisms, then we should observe functional interaction between them, that is, the overall effects should depend on the congruency of two colors in the voluntary and involuntary processes.

### Participants

Data from 4 participants were excluded from analysis because they had high proportion of equal (mixed) responses in at least one block. The remaining 7 participants had less than 3% equal response on average, and these equal responses were excluded from analysis.

### Procedure

Experiment 5 included three different tests: separate effect of mental imagery, separate effect of neon phantom color, and joint effect of mental imagery and neon phantom color within one trial. The first two isolated tests were identical to Experiment 4.

For the combined trials, participants were required to create a mental image of a given color for 4s, and passively view neon phantom color, both within the one trial. The order of imagined/neon phantom color was count-balanced within each block. The cue of which color to imagine was always right before the voluntary imagery period (Figure 3A).

**Figure 3.**
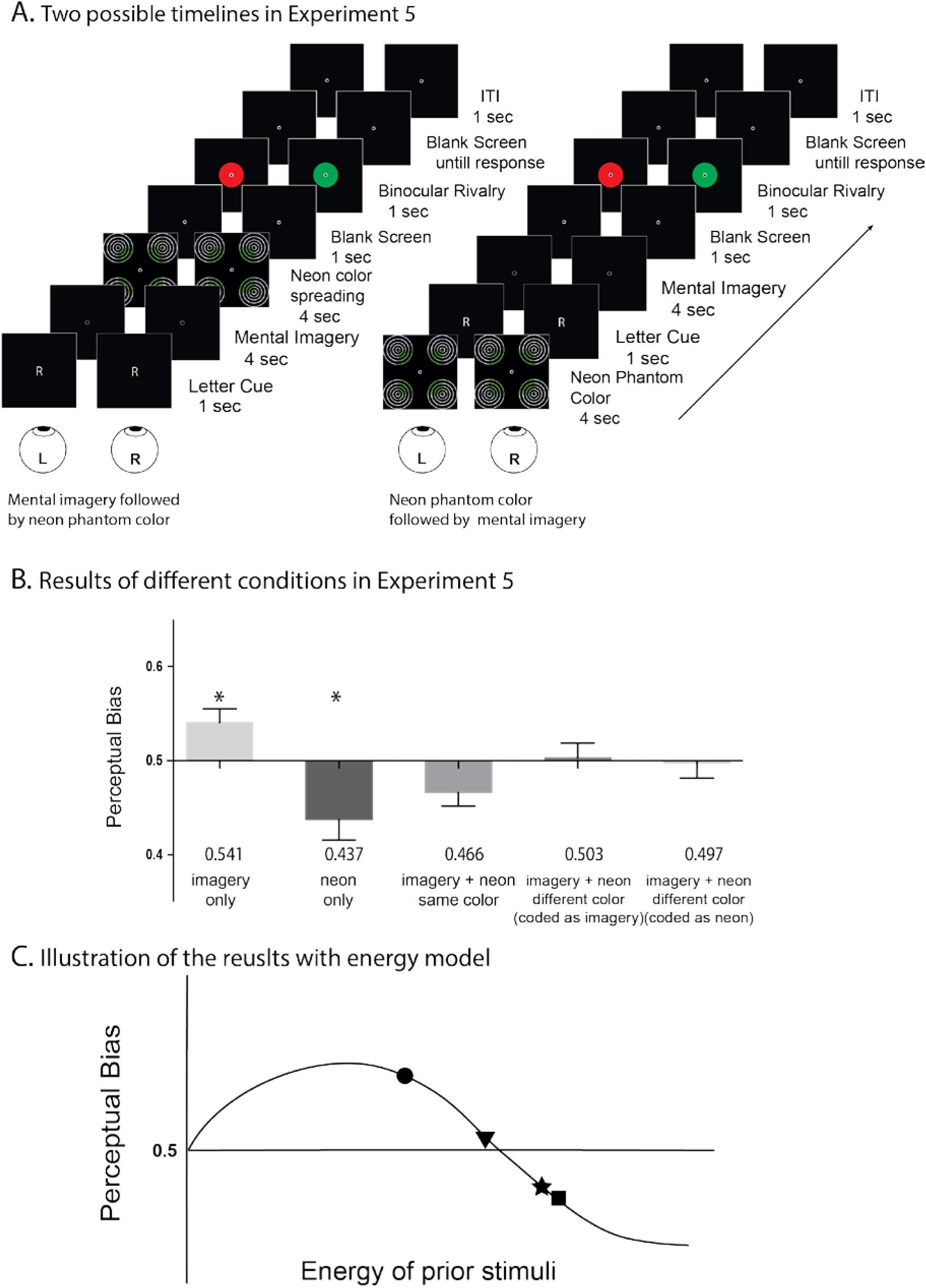
A. Timelines of Experiment 5 conditions. For half the trials, participants were instructed to do voluntary mental imagery first, for the other half, participants viewed neon phantom color first. Durations of the two color periods were both 4 seconds. B: Results of Experiment 5. Significant facilitatory effect of voluntary imagery and suppressive effect of neon phantom color were found. C: an illustration of results in Experiment 5 with Brascamp Model. Solid dot and square stand for effects of voluntary mental imagery and involuntary perceptual illusion. Solid five-point star stands for the joint effect of voluntary and involuntary phantom vision in the same color. Solid triangle stands for the location of effects when the two types of phantom vision were in different color. Explanation of this can be found in discussion session. Error bars stand for the SEM. * *p*<.05, ** *p*<.01

Mental imagery and neon phantom color viewing durations were both 4s. After the two tasks were completed, color rivalry was presented for 1s. Participants then pressed the relevant key to indicate the dominant color or a mix of the two. Time lines for the two types of trials can be found in Figure 3A.

Within one testing block, the imagined color and illusory color could be the same or different, and the number of trials for the two conditions were equal. There were 8 catch trials and 40 test trials in each block. The trials were randomly ordered. Each participant completed 3 blocks of each test.

### Results

In this experiment, for the separate effects, we found a significant facilitatory effect of color imagery on subsequent color rivalry perception (M = 0.541, t(6) = 2.83, *p* = .03, Cohen’s d = 2.3), and a significant suppressive effect of neon phantom color (M = 0.437, t(6) = 2.88, *p* = .028, Cohen’s d = 2.6). The results of color imagery replicated the previous studies (Chang et al., 2013; Pearson et al., 2008). The average decisional bias in catch trials in the imagery and neon color experiments were 53.0% and 46.7%, and neither of them was significantly different from chance level (*p*s > .06).

For the joint conditions, when imagery and neon phantom color were the same e.g. red+red, the overall effect on subsequent rivalry perception was suppressive (M = 0.466, t(6) = 2.4, *p* = .053). However, when the two color experiences were opposite (e.g. red+green), the effect on rivalry was not significantly different from chance (coded according to color of mental imagery: M = 0.503, t(6) = 0.175, *p* = .87).

We also analyzed the differences in perceptual biases across the three conditions, which were imagery only, imagery with same neon phantom color and imagery with opposite neon color. A repeated measure ANOVA showed that there was a significant difference across the three conditions (F = 6.84, *p* = .012).

These results can be taken to suggest that adding involuntary color can change the overall functional effect of voluntary phantom color, even when not occurring at the same time. When voluntary imagery was performed alone, it produced a priming effect on subsequent rivalry. When the same neon color was presented before or after voluntary color imagery, the overall effect was suppression. This indicates that the neon phantom color added to, or overpowered the imagery representation or memory trace produced by voluntary imagery. On the other hand, when neon phantom color was different from the imagined color, the overall sensory representation produced neither priming nor suppression. This indicates that the sensory representation formed by voluntary imagery could be weakened by subsequent involuntary phantom vision of a different color, which does not occur with the prior or subsequent presentation of a strong uniform yellow presented in a similar manner (Sherwood & Pearson, 2010).

## Experiment 6: The individual differences in phantom color interaction

In Experiment 5, 7 participants were tested, and we noticed possible individual differences among them. In the current experiment, we replicated this experiment with more participants.

### Participants

20 participants were recruited for this experiment, and all participants were included in the data analysis. The equal (mix) responses were 4% of all trials and they were removed from further analysis.

### Procedure

The experimental procedure was the same as Experiment 5. Each participant completed 2 blocks of separate color imagery test and neon phantom color test and 3 blocks of joint test.

### Results

For the separate tests, the perceptual bias from color imagery was M = 0.535, and it was not significantly different from chance (t(19) = 1.5, *p* = .15, Cohen’s d = 0.688). There was significant suppressive effect in neon phantom color test (M = 0.455, t(19) = 2.22, *p* = .039, Cohen’s d = 1.02). For the joint test, in the congruent condition, the perceptual bias was not significant from chance (M = 0.458, t(19) = 1.44, *p* = .165, Cohen’s d = 0.661). A significant facilitatory effect was found in the incongruent condition (M = 0.579, t(19) = 3.68, *p* = .002, Cohen’s d = 1.69). The average decisional bias in catch trials in the isolated color imagery, neon phantom color and joint effect experiments were 51.4%, 51.3% and 50.4%, respectively, and none were significantly different from chance level (*p*s > .19).

To examine the individual differences in the interaction between color imagery and neon phantom color, we checked the responses of each participant. First, the average responses of red dominance in rivalry perception accounted for 51% of all responses, and there is no significant difference between the proportions of red and green responses (t(38)=0.45, *p* = .66). This result indicates that participants did not have a pre-set color bias for rivalry.

Since the two phantom color processes were performed successively within each trial, there may be concerns that the “overall effects” were simply caused by the latter color and not the joint effect, that is, whether the order of the two processes influenced the perceptual biases. To exclude this factor, we analyzed whether there were differences in the order (see Figure 3A) for the four possible combinations between color imagery and neon phantom color (e.g., red neon phantom color followed by green color imagery and green color imagery followed by red neon phantom color). Results showed no significant difference between the opposite order configurations in the four conditions (*p*s > .2).

The above analyses suggest that the perceptual biases in joint condition are caused by the interaction between color imagery and neon phantom color. Based on this conclusion, we further explored the individual differences in the interaction between the two forms of phantom color. Here, we used the difference between biases in the congruent and incongruent condition as an index of functional interaction between color imagery and neon phantom color, since it reflects how much an individual sensory representation changes when the phantom color in the two processes are the same or not. This difference score can be thought of as a proxy measure of the interaction between the two different types of phantom color, and hence a proxy of overlap in their mechanisms. To illustrate, if there is no interaction between two forms of phantom color for one individual, then the perceptual biases should remain equal when we change the congruence of the two colors.

The histogram of differences between congruent and incongruent conditions is shown in Figure 4. We used sum of two Gaussian distribution model (GraphPad Prism, GraphPad Software, La Jolla California USA) to fit the data, and two Gaussian components were identified, with the means of the two Gaussians at −0.231 (SD = 0.04) and 0.017 (SD = 0.017), respectively (*R*^2^ = .539). When splitting all participants into two groups according to the identified Gaussian distributions, one group (with the mean at −0.231, interaction group) containing 9 participants and the other 11 (non-interaction group).

**Figure 4.**
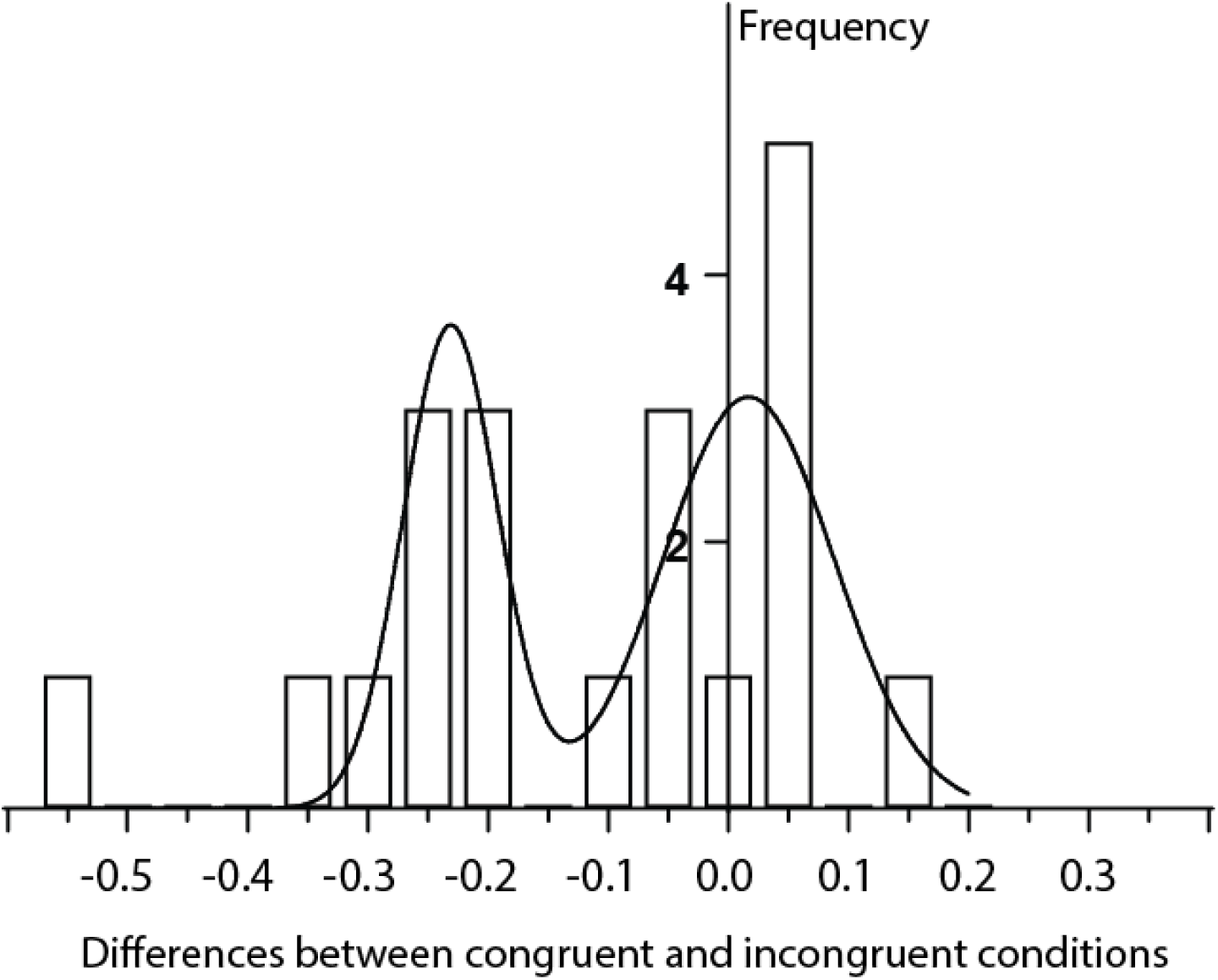
Distribution of differences between congruent and incongruent conditions in joint experiment in Experiment 6. The curved line illustrates two Gaussian distribution fitted to this data set.

This bimodal distribution suggests there might be two groups of individuals, those for whom the two types of phantom color interact, and those for whom they do not. This could be taken to suggest that there may be different neural mechanisms in phantom vision generation, and the degrees of similarity between voluntary and involuntary phantom vision vary across individuals.

## General discussion

Here we investigated the functional effects of involuntary neon phantom color on subsequent perception and how it relates to voluntary phantom color in the form of mental imagery. We found that passively experiencing neon phantom color led to suppressive effects on the subsequent binocular rivalry, and these effects were found in all six experiments. The suppression was attenuated when the color inducers were distorted. Perceptual factors such as concurrent background luminance and viewing durations could influence the suppressive effect, which was similar to weak perceptual stimuli and voluntarily generated mental stimuli (Chang & Pearson, 2017; Koenig-Robert & Pearson, 2016; Pearson et al., 2008), suggesting a sensory representation based neural mechanisms behind neon phantom color (Brascamp, Knapen, Kanai, van Ee, & van den Berg, 2007; Kanai & Verstraten, 2005; Pearson et al., 2008). The opposite biases induced by voluntary mental imagery and involuntary neon phantom color on subsequent binocular rivalry may suggest the sensory representations created by the two forms of phantom vision have different strengths or “energy”. According to the continuous model of facilitation and suppression (Brascamp et al., 2007), weak (low contrast or luminance) prior stimuli are more likely to induce facilitation, while strong or longer duration stimuli lead to suppression. Following this theory, the current results suggest that neon phantom color leads to relatively strong sensory representations, while imagery leads to weak sensory representation.

Besides the energy-based explanation, which claims a continuity between facilitatory and suppressive effects on binocular rivalry, there is another explanation that argues the opposite effects come from their different neural mechanisms. Neon phantom color may be the product of neural spiking in early visual cortex. Although we lack direct data in human subjects, single cell recordings in rats reported that after learning a motion sequence, neurons in rats V1 fired when only the starting point was shown, similar to when the complete motion sequence was presented (Xu, Jiang, Poo, & Dan, 2012). Neurons in monkey V1 corresponding to blind-spot showed significant activity when a bar moved across the blind-spot (Matsumoto & Komatsu, 2005). These data suggest that neon phantom color may be driven by neuronal spiking. On the other hand, for voluntary imagery, the feedback signals may play a role in suppressing the neural activation of colors other than the imagined one, in such a way that the imagined color primes. This modulatory effect of feedback from higher cortical areas would not lead to spiking in V1. This also explains why longer durations of imagery (up to 15 seconds), do not induce suppressive effects on subsequent rivalry, only greater levels of priming (Pearson et al., 2008).

The above two explanations, continuous energy and modulation vs. spiking, could also explain the results of the joint effects between imagery and neon phantom color. Using the continuous model (Brascamp et al., 2007), as shown in figure 3C, when mental imagery was added to the same passively viewed phantom color, an overall suppressive effect was observed (five-point star in Figure 3C). This suggests that the summed sensory representation was stronger than mental imagery alone (solid circle in Figure 3C). However, we also found that the joint effect of voluntary imagery and neon phantom color was not significantly stronger than the effect of neon phantom color alone, suggesting a non-linear addition. There are a few possible reasons for this. First, based on the Weber’s Law, the just noticeable difference is related to the absolute level of stimulation. Hence, the relatively low energy color imagery might have little effect on enhancing the overall behavioral response, when adding to the already high energy of the neon phantom color representation. An alternative may be that the mechanism for producing the sensory representation during voluntary imagery is largely shared by the neon phantom color process. In this case, under the neon phantom color only condition, the sensory representation of color experience may have already reached its peak, so adding imagined color could not make the overall sensory representation much stronger.

The modulation/spiking explanation of the joint effects experiment would suggest that imagery did not induce spiking, so the suppressive effect in the congruent condition solely came from the neon phantom color, which suggests that the joint effect is no different to the isolated neon phantom color. In the incongruent condition, the two processes may have cancelled each other out, thereby resulting in no effects on subsequent rivalry perception.

It seems that the current results can be explained by either of the two above theories. We may conclude that the correlation and interaction between the functional effects induced by the two forms of phantom vision rely on the neural activity in early visual areas that is largely influenced by feedback connections. However, how feedback signals shape activation in early visual areas during voluntary and involuntary phantom visual processes is largely unknown, and further research is needed.

Here we performed, to the best of our knowledge, the first study that directly compared voluntary and involuntary phantom perception, using a common indirect dependent measure: binocular rivalry. This quantitative measurement has been used in voluntary mental imagery studies and is regarded as an objective and reliable (Bergmann et al., 2015) technique to investigate visual mental imagery (for a review, see Pearson, 2014).

For research using the binocular rivalry paradigm, there are often concerns when dealing with subjective reports of stimuli. However, in the current study, we argue that the potential non-perceptual factors were excluded due to the following three reasons. First, catch trials that contained fake non-rivalrous stimuli were included in all experiments and our retained participants showed no bias. Second, no significant rivalry bias was found in the luminance condition in Experiment 2, suggesting that decisional or attentional effects, which should not be attenuated by background luminance, were not involved in the response. Third, both facilitatory and suppressive effects were found for mental imagery and neon phantom color, respectively. It would be unreasonable to assume a participant might have a prior decisional bias that flips direction between the two conditions.

Based on the results from current and previous work, we conclude that phantom color induced by neon color spreading can have a suppressive perceptual bias on subsequent binocular rivalry, and this effect can be strengthened by prolonged viewing and attenuated by background luminance. We also show that the effects produced by voluntary and involuntary phantom visual processes are related and can interact with each other over time. These results indicate that visual cortex is involved in involuntary phantom vision, and this type of sensory representation has similar characteristics to mental imagery. They may further suggest some shared mechanism between the processes of neon phantom color and mental imagery. These data also provide evidence in support of a common dichotomous framework for all phantom vision: voluntary and involuntary (Pearson & Westbrook, 2015), as has been proposed for visual attention (Connor, Egeth, & Yantis, 2004; Egeth & Yantis, 1997).

Further research can be conducted to answer remaining questions on voluntary and involuntary phantom vision. On the neural level, the origin of the feedback information in the neon phantom color and imagery is not clear, nor is the generation mechanisms of the two. On the behavioral level, whether color imagery can have suppressive effect and whether neon phantom color can have facilitatory effect on the subsequent binocular rivalry perception is worth investigating. The individual difference in the functional interaction between voluntary and involuntary phantom vison also needs further research. In the current study, only two forms of phantom vision were included, other forms of phantom vision should be studied in the future. We hope these suggested studies will provide more evidence to flesh out the dichotomous framework and the relationship between the two forms of phantom vision.

## Author contribution

All authors contributed to the development of the study concept and design. S. Chang performed the experiments, data collection, analysis, and interpretation under the supervision of J. Pearson. All authors drafted the manuscript and approved the final version of the manuscript for submission.

## Acknowledgement

We would like to thank Dr. Roger Koenig-Robert for providing comments on the manuscript.

